# Genomic changes associated with reproductive and migratory ecotypes in sockeye salmon (*Oncorhynchus nerka*)

**DOI:** 10.1101/117648

**Authors:** Andrew J Veale, Michael A Russello

## Abstract

Mechanisms underlying adaptive evolution can best be explored using paired populations displaying similar phenotypic divergence, illuminating the genomic changes associated with specific life history traits. Here we used paired migratory [anadromous vs. resident (kokanee)] and reproductive [shore- vs. stream-spawning] ecotypes of sockeye salmon (*Oncorhynchus nerka*) sampled from seven lakes and two rivers spanning three catchments (Columbia, Fraser, and Skeena) in British Columbia, Canada to investigate the patterns and processes underlying their divergence. Restriction-site associated DNA sequencing was used to genotype this sampling at 7,347 single nucleotide polymorphisms (SNPs), 334 of which were identified as outlier loci and candidates for divergent selection within at least one ecotype comparison. Eighty-six of these outliers were present in multiple comparisons, with thirty-three detected across multiple catchments. Of particular note, one locus was detected as the most significant outlier between shore and stream-spawning ecotypes in multiple comparisons and across catchments (Columbia, Fraser and Snake). We also detected several islands of divergence, some shared among comparisons, potentially showing linked signals of differential selection. The SNPs and genomic regions identified in our study offer a range of mechanistic hypotheses associated with the genetic basis of *O. nerka* life history variation and provide novel tools for informing fisheries management.

## Introduction

Understanding the mechanisms that facilitate local adaptation and identifying the loci underlying adaptive population divergence are important goals of evolutionary biology (Nielsen 2005; Nosil & Schluter 2011). The study of parallel phenotypic evolution, the repeated emergence of similar phenotypes linked to specific habitats, can provide important insights into these evolutionary processes (Arendt & Reznick 2008; Martin & Orgogozo 2013).

Parallel phenotypic evolution may be caused by the selection of identical adaptive mutations (Nachman *et al*. 2003; Paaby *et al*. 2010); however, there are also cases where shared phenotypes arise through different mutations at a given locus (Chan *et al*. 2010; Gross *et al*. 2009) or through mutations at entirely different underlying loci (Martin & Orgogozo 2013; Steiner *et al*. 2009). Where the same genetic changes are associated with parallel phenotypic evolution, there are multiple potential evolutionary scenarios that can account for this pattern (Welch & Jiggins 2014). One scenario is that the same adaptive alleles arise independently in each derived population – representing true molecular convergent evolution (Arendt & Reznick 2008). Alternatively, standing variation may exist in the ancestral population that is subsequently recurrently selected when populations colonize novel habitats (Barrett & Schluter 2008). Adaptive variation could also flow between ‘derived’ populations via the ancestral population – a situation known as the ‘transporter hypothesis’ (Bierne *et al*. 2013; Schluter & Conte 2009). There are many additional scenarios that can account for patterns of shared adaptations. The classic example of a genomic study of parallel evolution comes from the evolution of multiple freshwater populations of three-spined stickleback (*Gasterosteus aculeatus*), where anciently derived alleles in a gene responsible for armor reduction (Ectodysplasin-A) have been recurrently selected in multiple freshwater populations (Colosimo *et al*. 2005). This pattern has since been demonstrated across numerous other loci in stickleback (Jones *et al*. 2012). In this system, it appears likely that adaptive genetic variation that existed in one or more freshwater populations then spread to other freshwater populations via introgression with the marine population, although teasing apart the precise history of these events remains difficult (Welch & Jiggins 2014). Whether anciently derived, novel or introgressed from other populations, characterizing the genomic and geographical context of parallel phenotypic evolution provides vital clues about the mechanisms underlying adaptation.

Studies of the genomic bases of local adaptation have been facilitated by the advent of high-throughput genotyping methods, which allow for the identification and genotyping of thousands of genetic polymorphisms throughout the genome, enabling population genomic and association studies in non-model organisms (Baird *et al*. 2008; Davey *et al*. 2011; Miller *et al*. 2007). Studies that look to identify signals of adaptive divergence in wild populations often focus on sympatric populations that experience differential selective pressures, but are not highly diverged at neutral markers due to current or recent gene flow (Hemmer-Hansen *et al*. 2013; Roesti *et al*. 2015; Soria-Carrasco *et al*. 2014). This divergence with gene flow scenario has been hypothesized to generate ‘genomic islands of divergence’ displaying high genetic differentiation, contrasting with lower differentiation across the rest of the genome (Lotterhos & Whitlock 2015; Nosil *et al*. 2009; Via & West 2008). Although a suite of factors may influence the distribution and size of divergent regions including genetic conflict, mutation rates, genetic drift, and chromosomal structure, divergent selection and adaptation are often implicated (Bierne *et al*. 2011; Feder & Nosil 2010; Marques *et al*. 2016; Nosil *et al*. 2009). Genomic islands of divergence may arise because, while adaptive differences typically occur at a small number of loci, regions surrounding them are physically linked to the beneficial alleles (Kaplan *et al*. 1989; Maynard-Smith & Haigh 1974). This process can lead to regions of the genome containing multiple linked divergent single nucleotide polymorphisms (SNPs) around adaptive sites, a phenomenon termed divergence hitchhiking (Via 2012). These parts of the genome that underlie reproductive isolation or adaptation may then become resistant to the homogenizing effects of gene flow, and outlier detection tests can be used to identify these divergent SNPs (Foll & Gaggiotti 2008).

Salmonids are an exemplary taxonomic group to study the genetic basis of phenotypic and life history traits because abundant environmental variation, combined with precise natal homing, creates a situation ripe for differential local adaptation (Quinn 2005). Numerous examples of phenotypic variation and local adaptation have been reported within salmonid species (Fraser *et al*. 2011). These findings, along with their high cultural and economic value (Schindler *et al*. 2010), have contributed to salmonids being the primary taxonomic group where genomic research has been effectively applied to conservation and management (Garner *et al*. 2016; Shafer *et al*. 2015). The genomic bases for migration related traits have been a specific focus in salmon, with numerous quantitative trait loci (QTL) identified (Hecht *et al*. 2012; Norman *et al*. 2011). Genome-wide association studies (GWAS) have likewise detected several regions of the genome associated with migration (Hess *et al*. 2016). The genomic bases for other phenotypic traits including thermal tolerance, size and body condition have also been studied across a range of salmonids (Everett & Seeb 2014; Larson *et al*. 2016; Miller *et al*. 2012; Reid *et al*. 2005).

Sockeye salmon (*Oncorhynchus nerka*) exhibit tremendous life history and morphological variation, with several morphologically and ecologically divergent ecotypes linked to migratory and spawning behavior (Quinn 2005; Wood *et al*. 2008). All sockeye salmon spawn and spend their early life in freshwater, but while anadromous ecotypes later migrate out to sea, resident ecotypes (kokanee) remain in freshwater lakes throughout their life cycle (Groot & Margolis 1991; Quinn 2005). Sometimes these ecotypes occur sympatrically in the same lakes as genetically distinct populations (Foote *et al*. 1989; Taylor *et al*. 2000). It remains possible to re-anadromize kokanee by releasing them in rivers without access to lakes, but survival to maturation is lower by over an order of magnitude than similarly released anadromous sockeye salmon (Foerster 1947). Likewise, recently anthropogenically isolated anadromous sockeye salmon populations can survive and reproduce solely in freshwater, but these ‘residual’ sockeye salmon too have decreased survival and do not match aspects of the physiology, growth and morphology of kokanee (Smirnov 1959). Anadromous sockeye salmon and resident kokanee consistently exhibit a suite of heritable differences in size, morphology, early development rate, seawater adaptability, growth and maturation that appear to be divergent adaptations that have arisen from different selective regimes associated with their anadromous versus non-anadromous life histories (Gustafson *et al*. 1997). Kokanee populations are, however, polyphyletic having evolved from anadromous sockeye salmon through multiple independent postglacial freshwater colonization events (Taylor *et al*. 1996; Taylor *et al*. 1997;Wood *et al*. 2008). As the ‘kokanee phenotype’ is remarkably similar between catchments, these populations are examples of parallel convergent evolution across a wide range of traits (Taylor *et al*. 1996).

Both anadromous sockeye salmon and kokanee can be further subdivided into reproductive ecotypes, with each population exhibiting a specific spawning habitat preference. These include classical ‘stream (or river)-spawning’ ecotypes, ‘shore (or beach)-spawning’ ecotypes that spawn on the shallow submerged shorelines of lakes or island beaches, and ‘black-kokanee’ that also spawn on the lake benthos, but at depths down to 70m below the lake surface (Moreira & Taylor 2015). In some lakes, multiple reproductive ecotypes co-occur, while in others only one may be present. For kokanee, co-occurring reproductive ecotypes are generally morphologically indistinguishable, while for sockeye salmon populations, some (minor) phenotypic differentiation between adjacent reproductive ecotypes can occur (Quinn 2005). Survival of sockeye salmon eggs and fry is highly variable and influenced by a wide range of environmental variables (Whitlock *et al*. 2015). As the spawning habitats for each reproductive ecotype differ across many of these environmental factors, divergent natural selection is likely to play a role underlying ecotype divergence (Taylor *et al*. 2000).

Recently there have been several studies examining the genomic bases for ecotype divergence within sockeye salmon in single, but separate systems (Larson *et al*. 2017; Lemay & Russello 2015; Nichols *et al*. 2016). In Nichols *et al*. (2016), the genomic differentiation of population pairs of kokanee and anadromous sockeye salmon were evaluated from two lakes in the Snake River catchment. One of these pairs (Redfish Lake) also represented two different reproductive ecotypes (shore-spawning anadromous sockeye and stream-spawning kokanee). In Larson *et al*. (2017), several spawning sites representing different reproductive ecotypes of anadromous sockeye salmon (beach, river and stream) were sampled within a single catchment in Alaska. Likewise, Lemay & Russello (2015) investigated genomic differentiation between reproductive ecotypes (shore- vs. stream-spawning) of kokanee within Okanagan Lake, part of the Columbia River catchment. Although all three studies employed connectible restriction-site associated DNA sequencing (RADseq) methods (*SbfI;* Baird *et al*. 2008), their single-system scale largely precluded validation of the role of selection at detected outlier SNPs.

Here, we investigated the genomic patterns underlying ecotype divergence in sockeye salmon, employing RADseq of paired population samplings of migratory (anadromous versus resident) and reproductive (shore- versus stream-spawning kokanee) ecotypes sampled from seven lakes and two rivers spanning three catchments (Columbia, Fraser, and Skeena) in British Columbia, Canada. Our specific objectives were to: 1) reconstruct population genetic differentiation across a range of spatial scales including cross-drainage, among lake and within lake systems, including comparing our results to previously published studies; 2) identify outlier SNPs associated with ecotype divergence and investigate their generality as candidates for divergent selection; and 3) annotate surrounding genomic regions of the most predictive and divergent SNPs to inform mechanistic hypotheses related to *O. nerka* life history evolution.

## Materials and Methods

### Sampling

We used previously collected kokanee and sockeye salmon samples (either operculum punches or fin clips) or data from seven lakes and two rivers at the time of spawning (Table 1; Figure 1) across three historically-defined catchments in BC including: 1) Columbia River (Okanagan Lake, Wood Lake, Skaha Lake, Kootenay Lake, Okanagan River); 2) Fraser River (Anderson Lake, Seton Lake, Portage Creek); and 3) Skeena River (Tchesinkut Lake). Regarding the latter, although Tchesinkut Lake is currently connected to the Fraser River drainage, there is strong evidence that kokanee populations in this region were more closely associated with the Skeena River drainage historically (Taylor *et al*. 1996); for the purposes of this study, we will *a priori* follow the reconstruction of Taylor *et al*. (1996). In four of the lakes (Okanagan Lake, Wood Lake, Kootenay Lake, Tchesinkut Lake), stream-and shore-spawning kokanee co-occur, and were both sampled. Because these reproductive ecotypes are morphologically indistinguishable, all samples were obtained from spawning areas during the spawning period. In Kootenay Lake, we sampled both the North Arm and West Arm populations as previous studies have demonstrated significant spatial structure (Frazer & Russello 2013; Lemay & Russello 2012). Two lake/river systems have co-occurring kokanee and sockeye salmon, including one in the Columbia River drainage (Skaha Lake kokanee - Okanagan River anadromous sockeye salmon) and one in the Fraser River drainage (Anderson Lake/Seton Lake black kokanee - Portage Creek anadromous sockeye salmon). Portage Creek links Anderson and Seton Lakes, and has previously been demonstrated to be the most closely related anadromous sockeye salmon population to these kokanee populations (Moreira & Taylor 2015). Samples from Wood, Kootenay and Tchesinkut Lakes were originally collected for Frazer & Russello (2013). Skaha Lake and Okanagan River sockeye salmon samples were provided by the British Columbia Ministry of Forests, Lands and Natural Resource Operations and previously used in Veale & Russello (2016). Samples from Portage Creek, Anderson and Seton Lakes were originally collected for Moreira & Taylor (2015). Fully connectible data from Okanagan Lake were also used as previously collected in Lemay & Russello (2015).

**Figure 1.**
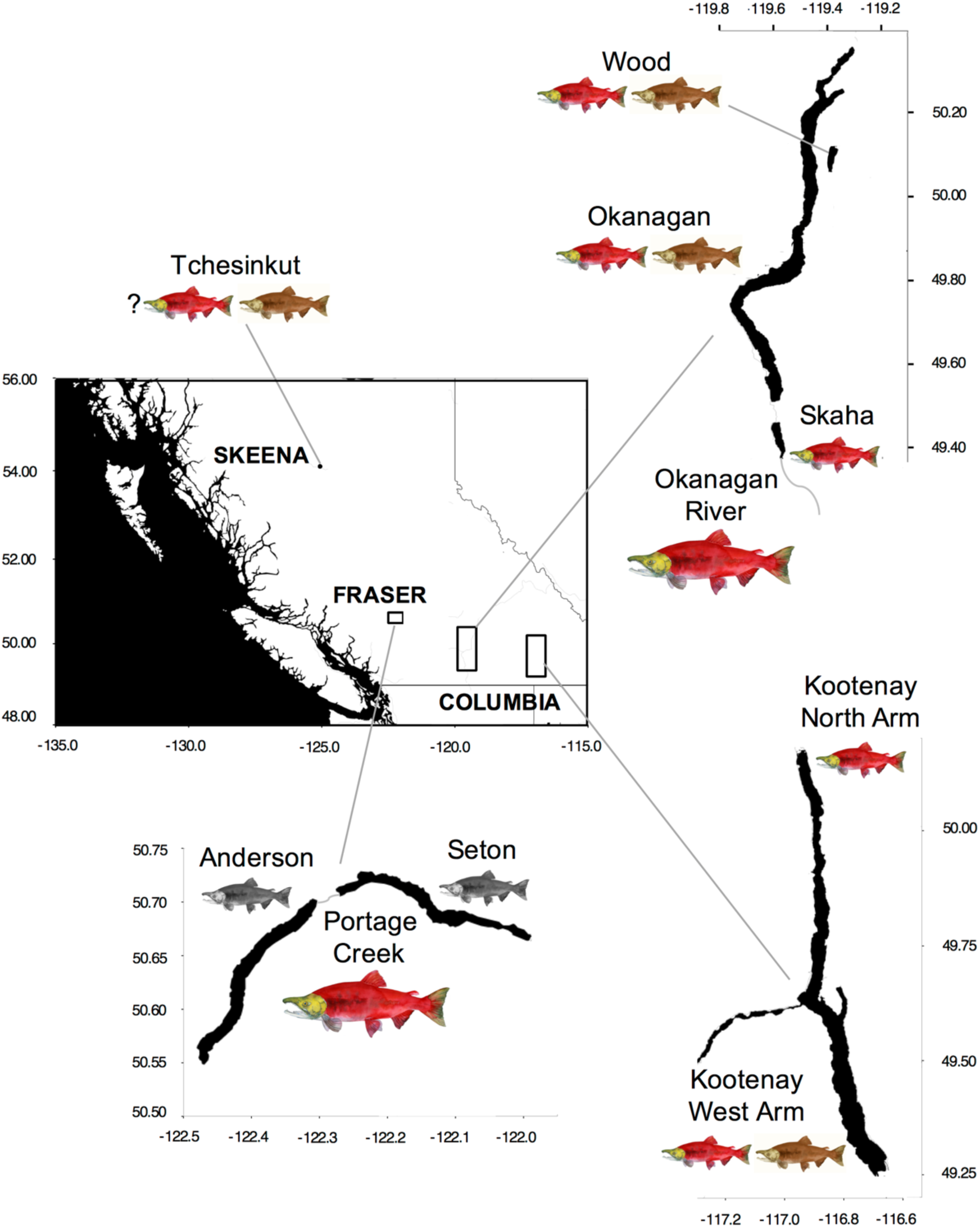
Map of British Columbia showing the four regions where kokanee and sockeye salmon samples were obtained. The historic river catchments are indicated in the inset. Size of *O. nerka* images denote migratory ecotype (large = anadromous sockeye salmon, small = kokanee), colors denote reproductive ecotype (red = stream-spawning, brown = shore-spawning, grey = black kokanee).

**Table 1.**
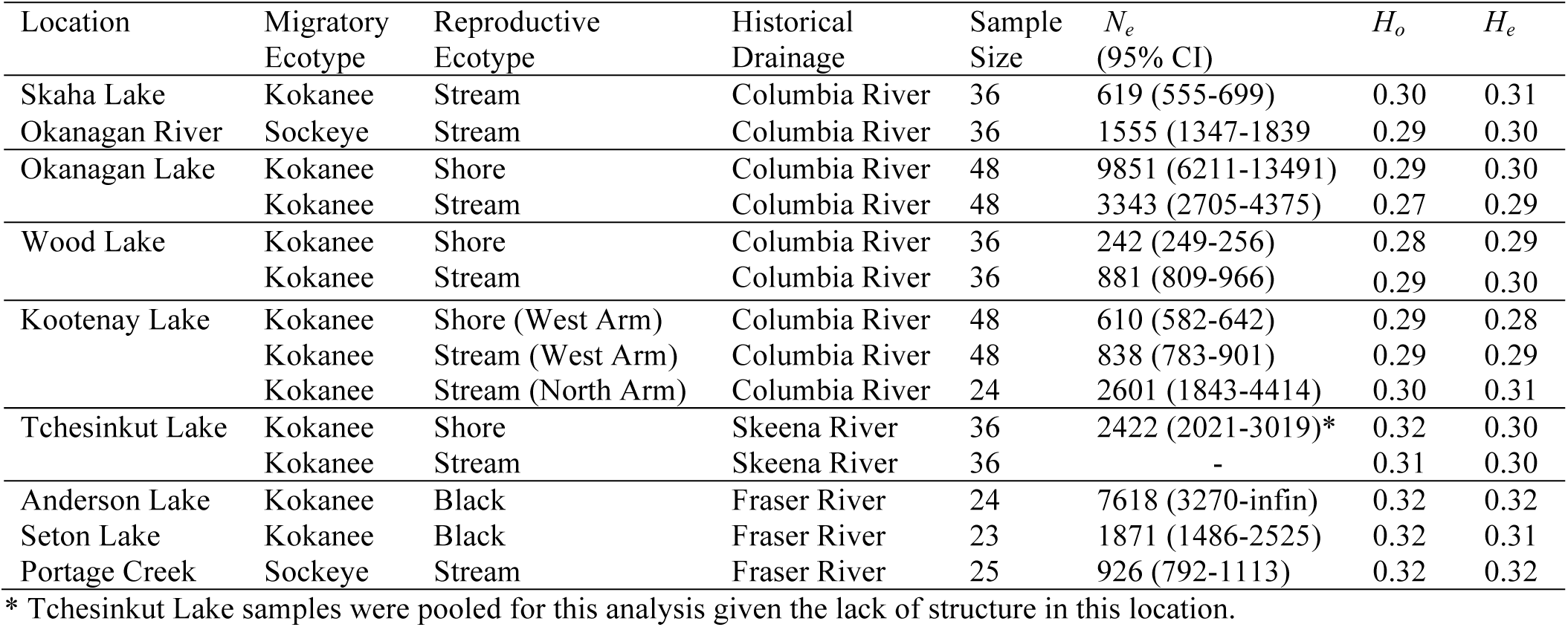
*Oncorhynchus nerka* (kokanee and sockeye salmon) samples used in this study, including migratory and reproductive ecotype designations, historical drainage, sample size, effective population size (*N_e_*), and heterozygosity [observed (*H_o_*) and expected (H_e_)].

| Location | Migratory Ecotype | Reproductive Ecotype | Historical Drainage | Sample Size | $N_e$ (95% CI) | $H_o$ | $H_e$ |
| --- | --- | --- | --- | --- | --- | --- | --- |
| Skaha Lake | Kokanee | Stream | Columbia River | 36 | 619 (555-699) | 0.30 | 0.31 |
| Okanagan River | Sockeye | Stream | Columbia River | 36 | 1555 (1347-1839) | 0.29 | 0.30 |
| Okanagan Lake | Kokanee | Shore | Columbia River | 48 | 9851 (6211-13491) | 0.29 | 0.30 |
|  | Kokanee | Stream | Columbia River | 48 | 3343 (2705-4375) | 0.27 | 0.29 |
| Wood Lake | Kokanee | Shore | Columbia River | 36 | 242 (249-256) | 0.28 | 0.29 |
|  | Kokanee | Stream | Columbia River | 36 | 881 (809-966) | 0.29 | 0.30 |
| Kootenay Lake | Kokanee | Shore (West Arm) | Columbia River | 48 | 610 (582-642) | 0.29 | 0.28 |
|  | Kokanee | Stream (West Arm) | Columbia River | 48 | 838 (783-901) | 0.29 | 0.29 |
|  | Kokanee | Stream (North Arm) | Columbia River | 24 | 2601 (1843-4414) | 0.30 | 0.31 |
| Tchesinkut Lake | Kokanee | Shore | Skeena River | 36 | 2422 (2021-3019)* | 0.32 | 0.30 |
|  | Kokanee | Stream | Skeena River | 36 | - | 0.31 | 0.30 |
| Anderson Lake | Kokanee | Black | Fraser River | 24 | 7618 (3270-infin) | 0.32 | 0.32 |
| Seton Lake | Kokanee | Black | Fraser River | 23 | 1871 (1486-2525) | 0.32 | 0.31 |
| Portage Creek | Sockeye | Stream | Fraser River | 25 | 926 (792-1113) | 0.32 | 0.32 |

\* Tchesinkut Lake samples were pooled for this analysis given the lack of structure in this location.

### RADseq genotyping

We constructed 14 novel RADseq libraries, each consisting of between 24 to 36 pooled, individually labeled *O. nerka* individuals (n = 150 total) following Baird *et al*. (2008) as modified in Lemay & Russello (2015). Genomic DNA was extracted using the NucleoSpin Tissue Kit (Macherey Nagel) following the manufacturer’s suggested protocol with the addition of RNase A (Qiagen). For each individual sample, 500 ng of DNA was digested using the *Sbf1* restriction enzyme (New England BioLabs Inc.). The barcodes used were six nucleotides in length, and each differed by at least two bases (Hohenlohe *et al*. 2010; Miller *et al*. 2012). During the library preparation, a sonicator (Bioruptor NGS; Diagenode) was used to shear DNA strands to a mean length of ∼500 base pairs, and a targeted fragment-size selection device (Pippin PrepTM; Sage Science) was used instead of gel extractions to isolate DNA fragments between 350 and 600 base pairs in length. One full lane each of Illumina HiSeq 2000 single end 110 bp sequencing was carried out for the 14 RADseq libraries at the McGill University and Génome Québec Innovation Centre, Montréal, Canada.

Previously collected RADseq data from identically prepared and sequenced Illumina libraries obtained from Okanagan Lake (Lemay & Russello 2015) were also included and reanalyzed from the raw data. All reads were trimmed, filtered and analyzed using the STACKS pipeline (Catchen *et al*. 2013) in order to create catalogues of comparable SNP loci. Initially, the PROCESS_RADTAGS module was used to separate reads by their barcode, remove low-quality reads (Phred quality score < 10), trim all reads to 100 base pairs in length, and remove any reads that did not contain the *Sbf1* recognition sequence. Next, the USTACKS module was used for the *de novo* assembly of raw reads into RAD tags. The minimum number of reads to create a stack was set at 3 (−m parameter in USTACKS), and the maximum number of pairwise differences between stacks was 2 (−M parameter in USTACKS). A catalogue of RAD tags was then generated using six *O. nerka* individuals from each putative population in CSTACKS. The distance allowed between catalogue loci (−n in CSTACKS) was increased to 1, after different trials were run to ensure loci were not inaccurately called as separate stacks. The execution of these components was accomplished using the STACKS denovo_map.pl script; in running this script, the optional −t flag was used to remove highly repetitive RAD tags during the USTACKS component of the pipeline. Following assembly and genotyping, the data were further filtered to maximize data quality. Using the POPULATIONS module, we retained only those loci that were genotyped in ≥80% of individuals and had a minor allele frequency ≥0.05 and a minimum stack depth of 10 (m in POPULATIONS) for each individual. Genotypic data were exported from STACKS in GENEPOP format (Raymond & Rousset 1995) and converted for subsequent analyses using PGD SPIDER v. 2 (Lischer & Excoffier 2012). Given the genome duplication event in the history of salmonid evolution, genomic samples are expected to contain a high proportion of paralogous sequence variants (PSVs); optimization of assembly parameters in salmonid systems is a fine balance between separating PSVs from their functional genes while not overly splitting informative variation. In general, we took an approach that was more likely to remove PSVs at the expense of potentially separating truly divergent sequence variants into separate loci. Following previous studies (Gagnaire *et al*. 2013; Hecht *et al*. 2012; Larson *et al*. 2017; Lemay & Russello 2015),we removed all loci that displayed significant deviation from Hardy–Weinberg equilibrium (HWE) as assessed using the method of Guo & Thompson (1992) implemented in the R package DIVERSITY (Keenan *et al*. 2013). Significance levels were adjusted for multiple comparisons using a 5% false discovery rate and the method of Benjamini & Yekutieli (2001). This analysis examined HWE in each of the 14 populations; to be removed from the data, a locus had to show significant deviations from HWE in at least three populations. This filtering provided a looser stringency for merging stacks, while still controlling for PSVs. For the outlier analyses, we retained multiple SNPs in each tag as per Nichols *et al* (2016) to ensure we did not lose potentially informative loci, however, for other analyses as noted below, we retained only the first SNP per tag to decrease linkage disequilibrium across samples.

RAD tags were mapped to the female sockeye salmon linkage map of Larson *et al*. (2016) and the RAD tags identified in sockeye salmon ecotypes in Nichols *et al*. (2016) and Larson *et al*. (2017) using BLAST-n as implemented in GENEIOUS (Drummond *et al*. 2006). We accepted any hit that was greater than E-25, and at least E-3 higher than any other alignment. See electronic supplementary materials for more details.

### Outlier locus detection and annotation

Due to the limitations of differentiation-based methods and the potentially high false positive rates when looking for outlier loci under divergent selection (De Mita *et al*. 2013; Vilas *et al*. 2012), we utilized two distinct approaches: 1) an *F_ST_* based outlier approach between *a priori* ecotype-pairs implemented in BAYESCAN (Foll & Gaggiotti 2008); and 2) a hierarchical Bayesian modeling approach implemented in PCADAPT (Luu *et al*. 2016). BAYESCAN was run first comparing ecotype paired populations (shore-and stream-spawning kokanee from within the same lake, and kokanee and anadromous sockeye salmon population pairs from the same catchment). Within Kootenay Lake this outlier detection was only carried out between shore- and stream-spawning populations in the West Arm of Kootenay Lake; North Arm individuals were excluded from this analysis, as spatial structure in this system has the potential to mask detection of candidate loci. In addition, given the low neutral divergence between Okanagan Lake shore- and stream-spawning kokanee, Kootenay Lake West Arm shore- and stream-spawning kokanee, and Anderson and Seton Lakes kokanee (*F_ST_* ≈0.01; see results), we treated each of these as single lake populations for the purposes of *F_ST_* outlier tests between kokanee and sockeye salmon pairs. We also used BAYESCAN to evaluate combined ecotype datasets: 1) all shore-spawning kokanee vs. all stream-spawning kokanee, and 2) all Okanagan catchment kokanee (Okanagan Lake, Wood Lake, Skaha Lake) vs. Okanagan river anadromous sockeye salmon. For each analysis, BAYESCAN was run using 10,000 output iterations, a thinning interval of 10, 20 pilot runs of length 10,000, and a burnin period of 10,000, with prior odds of the neutral model of 10. We recorded all loci with a q-value of 0.2 or less - which equates to a false discovery rate of 20% (note: q-values are much more stringent than p-values in classical statistics).

We also conducted hierarchical outlier detection on the complete dataset and on reduced subsets of the data (within each lake and/or catchment) as implemented in PCADAPT (Luu *et al*. 2016). The number of Principle Components retained (K) for each analysis was determined by the graphical approach based on the scree plot (Jackson 1993) as recommended by Luu *et al*. (2016). We recorded all loci with a q-value of 0.2 or less, and identified the principle component associated with each outlier. Within each lake-level analysis, we also identified potentially misidentified individuals using the clustering output.

In addition to mapping to the female sockeye salmon linkage map (Larson *et al*. 2016) and the sockeye salmon RAD tags (Larson *et al*. 2017; Nichols *et al*. 2016) described above, all loci identified as outliers in either analysis were compared with published loci on Genbank using BLAST-n in order to identify nearby genes. We also used BLAST-n specifically in the *Salmo salar* and *O. mykiss* genomes to map each outlier locus, accepting any hit that was greater than E-25, and at least E-3 higher than any other alignment. See electronic supplementary materials for more details.

### Island of divergence mapping

We used a Gaussian kernel smoothing technique (Hohenlohe *et al*. 2010; Larson *et al*. 2017; Larson *et al*. 2014b) to identify putative islands of divergence based on the *F_ST_* of loci that could be placed on the sockeye salmon linkage map of Larson *et al*. (2016). We used a window size of 5 cM and a stepwise shift of 1 cM for this analysis, and *F_ST_* values were weighted according to their window position as described by Gagnaire *et al*. (2013). These values were selected based on those used by Larson *et al*. (2017), which had been optimized for this particular linkage map. We identified highly differentiated windows by randomly sampling *N* loci from the genome, where *N* was the number of loci in the window, and comparing the average *F_ST_* of those loci to the average *F_ST_* of the loci in the window. This sampling routine was conducted 1,000 times for each window.

If a window exceeded the 90% percentile of the sampling distribution, the number of bootstrap replicates was increased to 10,000. Contiguous windows that contained at least two loci and exceed the 99% percentile of the distribution after 10,000 bootstrap replicates were classified as islands of divergence. Along with the formal tests for islands of divergence, we compared the locations of outlier loci identified in different pairwise tests to see if there were patterns of shared genomic regional divergence beyond the sharing of the same SNP.

### Demography, diversity and population structure

We calculated locus- and population-specific estimates of observed (*H_o_*) and expected heterozygosity (*H_e_*) using the neutral-only dataset (all outliers removed, one SNP per retained RAD tag) as implemented in ARLEQUIN (Excoffier & Lischer 2010). Effective population sizes were estimated using the linkage disequilibrium method implemented in NEESTIMATOR (Do *et al*. 2014).

We investigated the number of populations (or clusters) represented in our entire sampling using FASTSTRUCTURE (Raj *et al*. 2014) and the neutral SNP dataset, default parameters, a logistic prior, and *K* from 1 to 16. The appropriate number of model components that explained structure in the dataset was determined using the *chooseK.py* function (Raj *et al*. 2014). In order to test for unrecognized substructure in the global FASTSTRUCTURE analysis, we grouped populations according to their cluster membership and repeated the above analyses on the reduced datasets as recommended by Pritchard *et al*. (2000). Results for the identified optimal values of *K* were visualized using DISTRUCT (Rosenberg 2004). We further calculated both locus-specific and population-wide *F_ST_* (Weir & Cockerham 1984) using DIVERSITY (Keenan *et al*. 2013), and reconstructed a two-dimensional NeighbourNet network (Bryant & Moulton 2004) based on the pairwise neutral-only population *F_ST_* values as implemented in SPLITSTREE 4.0 (Huson & Bryant 2006). See electronic supplementary materials for more details.

## Results

### RADseq genotypic data and alignment

Following RAD sequencing, processing and filtering, we collected genotypic data at 8,436 SNPs (30X average coverage) for 504 individuals representing anadromous sockeye salmon and resident kokanee ecotypes from seven lakes and two rivers across three historically-defined catchments in BC (Figure 1, Table 1). After filtering for within population deviations from Hardy-Weinberg equilibrium, this number was reduced to 7,347 SNPs across 6,568 RAD tags. Of these tags, 41 (0.62%) contained three SNPs per tag, 738 (11.2%) contained two SNPs per tag, while 5871 (89.4%) contained only one SNP.We removed 32 individuals due to low coverage (<50% SNPs retained), giving a final dataset of 472 individuals across the 14 sampling units (Table 1).

Of the 6,568 retained tags in our study, 43% (2,796) were unambiguously mapped to the sockeye salmon linkage map of Larson *et al*. (2016). Furthermore, 1,714 (26%) of our tags matched those retained by Nichols *et al*. (2016), equating to 66% of the 2,593 RAD tags used in their study.

### Demography, diversity and population structure

Estimates of population-level observed and expected heterozygosity based on 6,234 neutral loci [outliers removed (see below); one SNP per RAD tag] were virtually identical across all ecotypes and locations (0.29-0.32; Table 1). Effective population sizes varied by an order of magnitude between populations (242-9851; Table 1), with no clear trends observed by spawning location or ecotype (Table 1).

The FASTSTRUCTURE analysis based on the neutral dataset provided evidence for eight distinct clusters in our data, grouping together Anderson and Seton Lake black kokanee, Kootenay West Arm shore-and stream-spawning kokanee, Okanagan Lake shore- and stream-spawning kokanee combined with Skaha Lake kokanee, and Tchesinkut Lake shore- and stream-spawning kokanee (Figure 2). In this global analysis, Wood Lake included a large contribution from Okanagan/Skaha Lake, with Skaha Lake also showing admixture with Okanagan River sockeye salmon. The presence of substructure in the data was further explored in each of these systems individually; only Wood Lake revealed two distinct clusters corresponding to shore-and stream-spawning kokanee when analyzed independently as well as a part of an Okanagan Basin only analysis (data not shown).

The phylogenetic network based on the neutral dataset revealed a clear separation between the historical Skeena River drainage site (Tchesinkut Lake) and all the Columbia-Fraser drainage lakes and rivers (*F_ST_*=0.37−0.45) (Table S1, Figure S1). Kootenay Lake kokanee exhibited significant population structure between the North Arm and West Arm populations (*F_ST_*=0.13), which were both more closely related to Fraser River drainage sites (Anderson Lake/Seton Lake kokanee and Portage Creek sockeye salmon) than to the Columbia River drainage sites sampled in the Okanagan Basin (Okanagan, Wood and Skaha Lakes). At a finer level, shore-and stream-spawning populations sampled in the same lake (Okanagan, Wood, Kootenay, Tchesinkut) were more closely related to each other than to corresponding ecotypes in other lakes without exception (Figure S1). These paired kokanee ecotype samplings exhibited widely differing levels of intra-lake divergence and in some cases, possible introgression. Wood Lake shore- and stream-spawning kokanee were moderately divergent (*F_ST_*=0.060), while Okanagan Lake shore-and stream-spawning kokanee and Kootenay Lake West Arm shore- and stream-spawning kokanee were considerably less so (*F_ST_*=0.008; Table S1, Figure S1). There was minimal differentiation between kokanee sampled on the shore and at the mouth of the stream in Tchesinkut Lake (*F_ST_*=0.003; Table S1, Figure S1).

The PCADAPT analyses both highlight population structure and simultaneously identify outlier loci associated with this structure. Here, we will present the population structure indicated by these analyses (see below for outlier analysis). We found the optimal number of principle components to be nine for the full dataset. For most of the principle components retained, geographic structure was revealed rather than ecotype differentiation (Figure S2). The first two principle-components match the structure indicated by the phylogenetic network - PC1 separating Tchesinkut Lake kokanee from all other populations, PC2 separating Kootenay Lake kokanee with all other populations (Figure S2A). PC3 separates the current Fraser River populations (Anderson & Seton Lake Kokanee, Portage Creek anadromous sockeye salmon) from the other populations (Figure S2B). PC4 is more complex, with populations spread out, though with a less clear pattern (Figure S2B). There is some separation of shore/stream-spawning populations on this axis including: Wood Lake shore- and stream-spawning kokanee, Anderson/Seton Lake shore-spawning kokanee, and Okanagan River stream-spawning sockeye and Skaha Lake stream-spawning kokanee; however, other shore/stream-spawning population pairs are not separated. PC6, PC7, PC8, and PC9 each separated one population from all of the others (Kootenay North Arm kokanee, Portage Creek sockeye salmon, Wood Lake stream-spawning kokanee, and Skaha kokanee, respectively; Figure S2C-E).

The PCADAPT results were more interpretable relating to ecotype divergence when conducting analyses at the lake-level. Within the Anderson/Seton Lake and Portage Creek system, two principle components were optimal, with PC1 separating Portage Creek anadromous sockeye salmon from the two kokanee populations, and PC2 separating Anderson and Seton Lake shore-spawning kokanee from each other (Figure S3A). Similarly within Kootenay Lake, two principle components were optimal, with PC1 separating the North Arm stream-spawning kokanee, from the two West Arm populations, and PC2 separating the West Arm stream-spawning kokanee from the West Arm shore-spawning kokanee (Figure S3B). Three potentially misidentified individuals were detected between the two West Arm kokanee populations. Within the Wood Lake system, two principle components were also optimal, with PC1 separating the shore- and stream-spawning kokanee populations, and PC2 separating off two shore-spawning individuals (Figure S3C). Again, some potentially misidentified individuals were detected – two shore-spawners in the stream-spawning cluster. For the three remaining within-lake comparisons, it was unclear if one or two principle components were optimal, so while we display two principle components for each of these analyses, we acknowledge that potentially only the first is supported. From the Okanagan Lake analysis, PC1 separates perfectly the shore- and stream-spawning kokanee (Figure S3D). We further annotated the PC plot with the various spawning locations sampled in case there was a spatial pattern to PC2, however, all streams and shore-spawning locations were mixed evenly along this axis, showing no detectable within-lake spatial structure beyond ecotype differentiation. From the Tchesinkut Lake analysis, no ecotype separation was evident in any principle component, with complete overlap between ‘ecotypes’ (Figure S3E). Within the Skaha Lake kokanee/Okanagan River anadromous sockeye analysis, PC1 fully separated each ecotype (Figure S3F).

**Figure 2.**
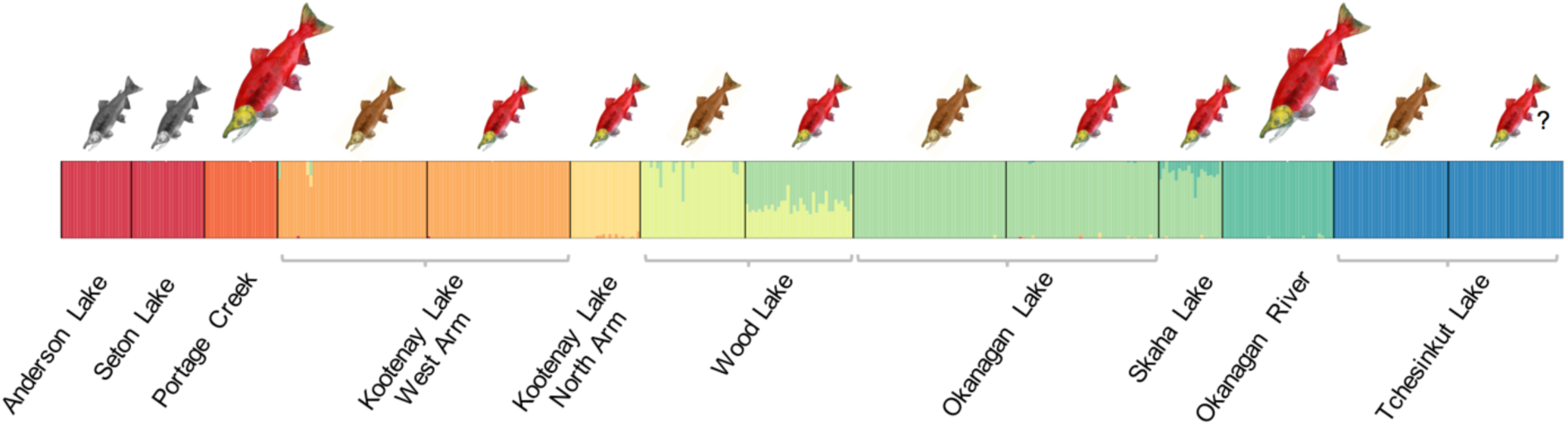
FASTSTRUCTURE (Raj *et al*. 2014) plot based on genotypic data at 6,234 neutral SNPs showing the proportion of cluster membership at K=8. Vertical bars indicate individuals, colors represent proportion of cluster membership. Size and color of *O. nerka* images as in Figure 1.

### Outlier SNP analyses

The two outlier analyses methods used evaluated different kinds of divergence – pairwise divergence between *a priori* groups in BAYESCAN, and *a posteriori* evaluations of divergence between computed clusters in PCADAPT. These analyses are therefore sometimes comparable when these two groupings matched, but incomparable (though complementary) when the principle components did not match the predefined populations. These complementary analyses therefore add different views of outlier loci and their underlying drivers.

Because the principle components identified by PCADAPT in the complete dataset analyses either primarily reflected geographic differentiation (PCs 1-3) or single-population-specific outliers (PCs 6 – 9), we report these outlier SNPs (Table S2), but do not analyze them in detail here. Rather, we focus on the genomic changes underlying ecotype divergence that were identified by both outlier detection approaches (BAYESCAN and PCADAPT).

Numerous outlier loci were detected across the three primary pairwise ecotype comparisons including Anderson/Seton Lake shore-spawning kokanee vs. Portage Creek anadromous sockeye salmon (n = 68), Okanagan Basin kokanee (Okanagan Lake, Wood Lake, Skaha Lake) vs. Okanagan River anadromous sockeye salmon (n = 34), and all shore-spawning kokanee vs. all stream-spawning kokanee (n = 32; Figure 3). Of these, three outliers (R40949, R68810, R122729) were shared in multiple comparisons. In particular, R68810 was identified in all three comparisons, and was notably the most significantly divergent in the shore-spawning vs. stream-spawning kokanee comparison by several orders of magnitude (Figure 3).

Across these three combined comparisons, along with the four-paired shore/stream spawning kokanee comparison within lakes, and the six-paired kokanee/anadromous sockeye salmon comparisons, a total of 334 outlier loci were detected across the two methods. Of these, 86 were detected in multiple comparisons, including 11 that were detected as outliers between sockeye salmon ecotypes in Nichols *et al*. (2016), and 6 that were recorded as outliers between reproductive ecotypes of anadromous sockeye salmon in Larson *et al*. (2017) (Table 2, Table S3). Comparing the outliers detected in BAYESCAN with those in directly comparable PCADAPT analyses, 37% were identified by both analyses, 37% were identified by BAYESCAN only, and 26% were identified by PCADAPT only (Table S3).

Shared outlier loci between pairwise comparisons were most common between Okanagan River anadromous sockeye salmon in comparisons with nearby lakes within the Okanagan Basin (Okanagan Lake, Wood Lake and Skaha Lake) as well as between the two populations in Kootenay Lake. Seven of the outlier loci detected between Anderson and Seton Lake kokanee and Portage Creek anadromous sockeye salmon were also detected in other population comparisons, six of which have high confidence annotations (Table 3).

The common shore/stream outlier locus R68810 was independently detected in three paired kokanee ecotype comparisons (Tables S3, S4) and in Nichols *et al*. (2016) (Table 2). This shared outlier locus had the highest differentiation in both Anderson-Seton Lakes kokanee/Portage Creek sockeye salmon (*F_ST_* = 0.83), and in Okanagan Lake kokanee shore- and stream-spawning comparisons (*F_ST_* = 0.92), and was significantly differentiated between Wood Lake kokanee shore- and stream-spawning populations (*F_ST_* = 0.43). R68810 was also detected as a highly significant outlier between two other shore-spawning kokanee and stream-spawning sockeye salmon pairs [Okanagan Lake shore-spawning kokanee/Okanagan River sockeye (*F_ST_* = 0.97), and Wood Lake shore-spawning kokanee/Okanagan River sockeye salmon (*F_ST_* = 0.90); Table S4].

**Figure 3.**
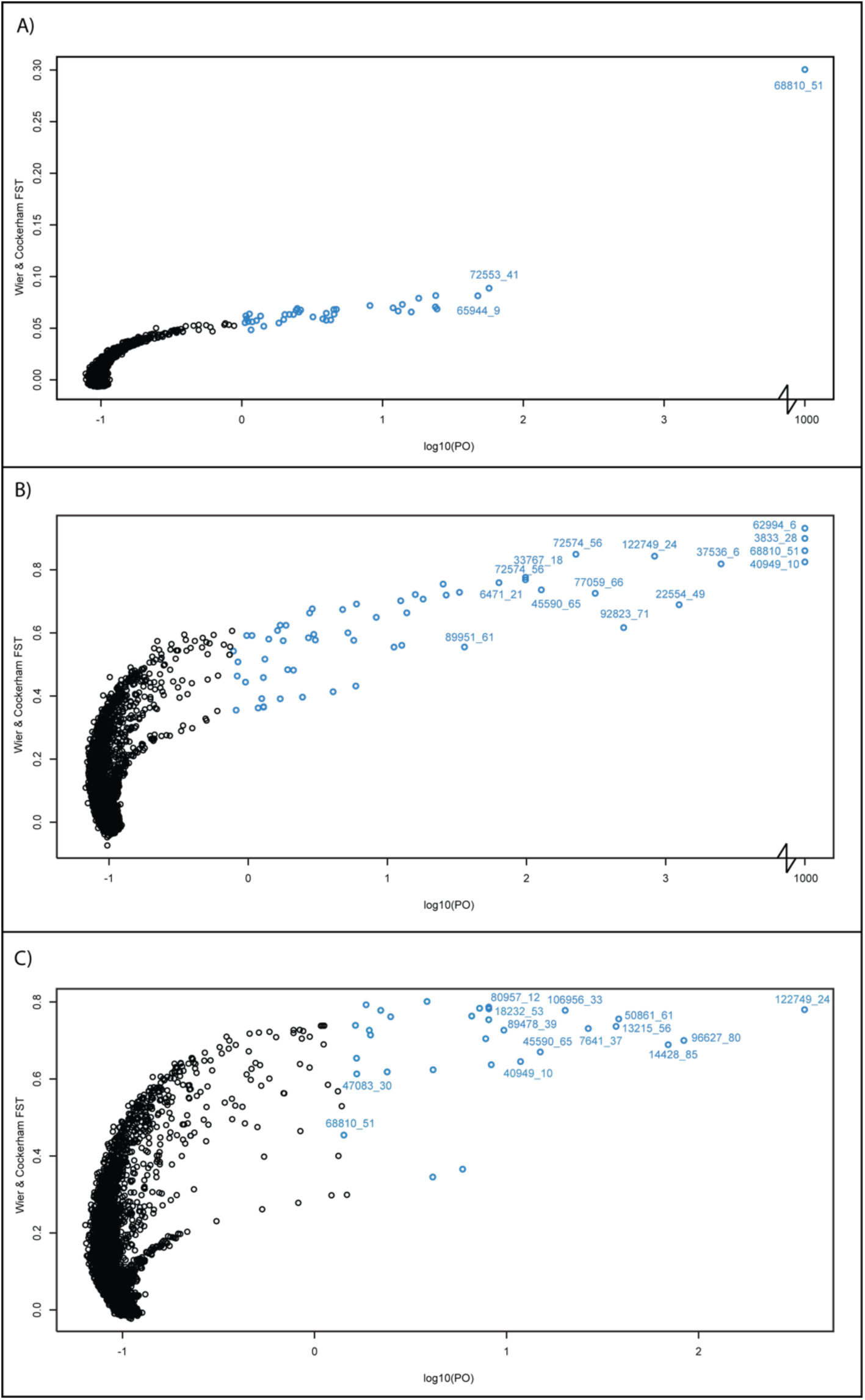
*F_ST_* outlier plots for the three primary combined ecotype comparisons performed in BAYESCAN (Foll & Gaggiotti 2008) including: A) shore- versus stream-spawning kokanee; B) Anderson/Seton Lake shore-spawning kokanee versus Portage Creek anadromous sockeye salmon; and C) Okanagan Basin lakes kokanee versus Okanagan River anadromous sockeye salmon. Note the disjoint x-axis, with loci that had a q-value of 0.0000, and a Log10(PO) of 1000 brought in to aid visibility.

**Table 2.** Outlier loci independently detected in three studies of anadromous sockeye salmon and/or kokanee across multiple sites and catchments. Color shading indicates the comparison within which an outlier was detected including: shore/stream-spawning (light gray), kokanee/anadromous sockeye salmon (dark gray), and mixed shore/stream-spawning and kokanee/anadromous sockeye salmon comparisons (gray grid). Study-specific RAD tag codes are also indicated.

| Site | Multiple Sites | Redfish Lake | Alturas Lake | Bristol Bay |
| --- | --- | --- | --- | --- |
| Catchment | Columbia, Fraser, Skeena Rivers<br>(British Columbia, Canada) | Snake River<br>(Idaho, USA) | Snake River<br>(Idaho, USA) | Wood River<br>(Alaska, USA) |
| Reference | This Study | Nichols *et al.* (2016) | Nichols *et al.* (2016) | Larson *et al.* (2017) |
| RAD tag | 24343_34 | 64477 | 64477 | 41305 |
|  | 27613_58 | 67582 | - | - |
|  | 42694_49 | - | - | 86418 |
|  | 54987_82 | 10067 | - | - |
|  | 58500_65 | - | - | 11863 |
|  | 59110_15 | 37622 | - | - |
|  | 62994_06 | 84113 | - | - |
|  | 68420_46 | - | - | 82702 |
|  | 68810_51 | 57884 | - | - |
|  | 84183_30 | - | 30017 | - |
|  | 86537_58 | - | 22082 | - |
|  | 89327_44 | - | - | 88727 |
|  | 93027_29 | - | - | 12430 |
|  | 105177_36 | - | 13591 | - |
|  | 114019_82 | 24175 | 24175 | - |
|  | 119684_49 | 20138 | - | - |

### Islands of Divergence

In all three of the primary comparisons, we found regions of the genome with elevated divergence (Figure 4, Table S5). Due to the relatively low coverage of SNPs across this linkage map (38% of SNPs retained in our study mapped), there may be other regions with shared elevated divergence that could not reach statistical significance given the sampling level.

Of particular note are putative islands of divergence shared between our comparisons and/or with other studies. Islands LG12_3 and LG12_4 were almost identical in location, with LG12_3 identified between the Anderson and Seton Lake shore-spawning kokanee and the Portage Creek anadromous sockeye salmon, while LG12_4 was identified in the combined paired kokanee shore/stream comparison. Similarly, LG24_1, LG24_2 and LG24_3 have overlapping regions and were identified in both the Anderson and Seton Lake shore-spawning kokanee/Portage Creek anadromous sockeye salmon comparison, and the Okanagan Basin Lakes kokanee/Okanagan River anadromous sockeye salmon comparison.

Comparing our results with those of Larson *et al*. (2017), we also found two shared islands of divergence. Our LG12_1 and LG12_2 overlap LG12_1 in Larson *et al*. (2017), one of the most prominent island of divergence between their beach-spawning and river-spawning sockeye salmon. Finally our LG15_1 matches LG15_1 in Larson *et al*. (2017), a region containing the known MHC genes.

**Figure 4.**
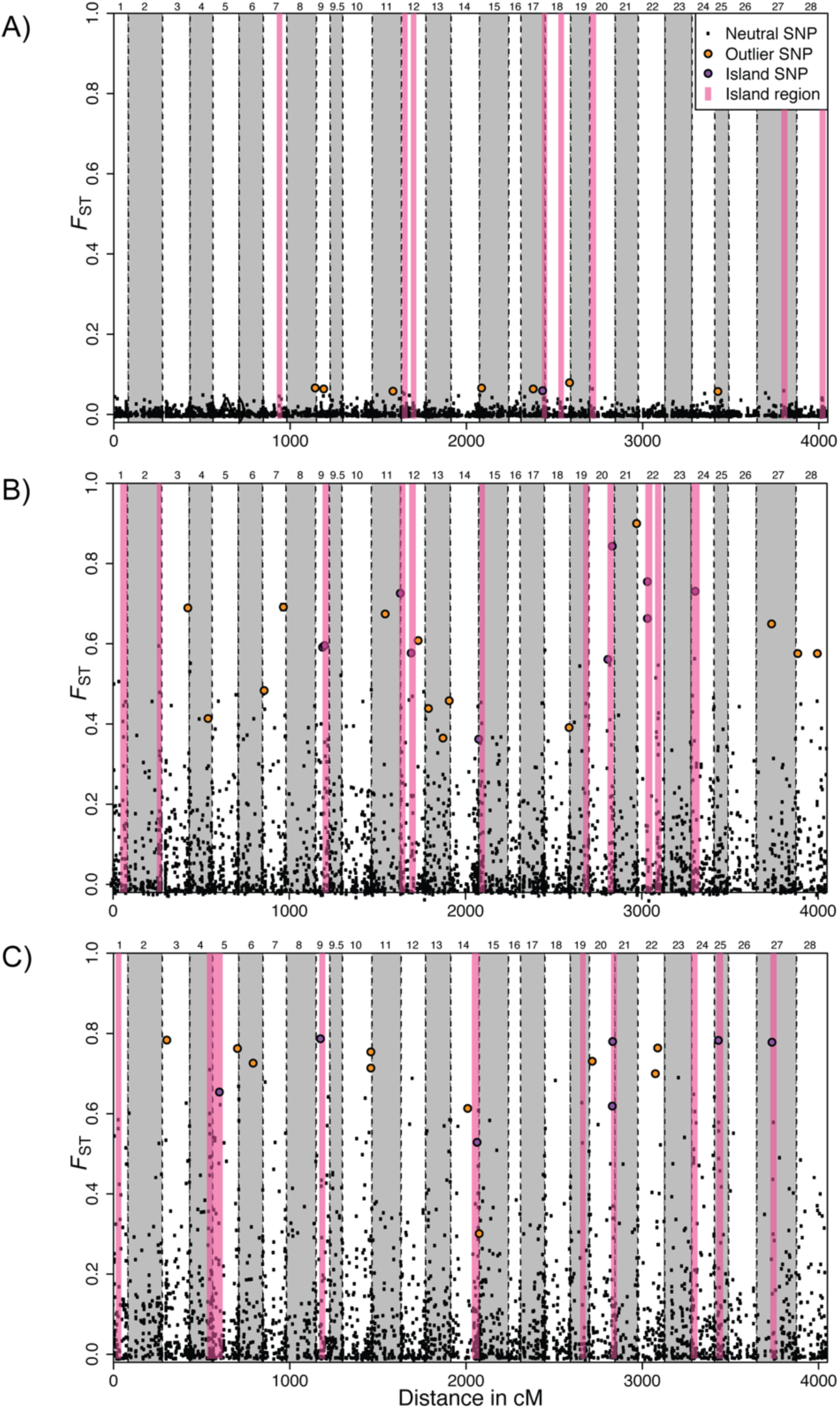
Plots of genetic differentiation (*F_ST_*) across each linkage group in the sockeye salmon linkage map (Larson *et al*. 2016) for the three primary combined ecotype comparisons including: A) shore-versus stream-spawning kokanee; B) Anderson/Seton Lake shore-spawning kokanee versus Portage Creek anadromous sockeye; and C) Okanagan Basin lakes kokanee versus Okanagan River anadromous sockeye. Islands of divergence identified using a kernel smoothing technique are denoted as “Island regions”. The “Outlier SNP” designation denotes outlier loci that are found outside of islands of divergence, and the “Island SNP” designation indicates outlier loci that are found within islands of divergence.

### Alignment and annotation of outlier loci

One hundred and twenty-seven outlier loci unambiguously aligned within the Atlantic salmon (*Salmo salar*) genome, including the common shore-stream outlier R68810 (Table S4). In addition to the specific annotations of these individual loci detailed in Table S4, we detected three instances where two RAD tags overlapped by four base pairs (R50861/R89478, R95009/R97203, R88765/R17286) flanking the same restriction site. These RAD tag pairs (and the associated outlier SNPs) therefore should each be treated as single locus. More distant potential linkage was also detected based on the alignment with the *S. salar* genome, with R40949 and R74283 only 367 kilobase pairs away from each other in the *S. salar* genome. These two loci were among the most commonly identified sockeye/kokanee outlier loci in our comparisons (4 comparisons and 5 comparisons, respectively), suggesting they are potentially linked to the same genomic region under differential selection, and within the same island of divergence - though precisely where on the sockeye salmon linkage groups remains uncertain as neither mapped to the linkage groups of Larson *et al*. (2016).

## Discussion

Our results offer important insights into the evolutionary history of *O. nerka*, inferring genomic mechanisms underlying adaptive divergence and identifying specific genes that exhibit parallel patterns across multiple ecotype-pairs sampled from different catchments. Furthermore, the connectibility of our genotypic data with those from previously published studies employing *SbfI* RADseq enable us to explore the generality of these mechanisms and loci at a broader scale.

### Evolutionary history of sockeye salmon

Although our study was not designed to provide a comprehensive investigation of *O. nerka* evolutionary history in BC, the patterns of divergence we observed between sampled populations based on 6,234 neutral SNPs reinforce findings from previous studies (Taylor *et al*. 1996). We found a clear separation between the historical Skeena River drainage population sampled in Tchesinkut Lake and all kokanee and sockeye salmon populations sampled from the Columbia-Fraser River drainages, consistent with the multiple glacial refugia hypothesis set forth by Taylor *et al* (1996). Within the Fraser River drainage, Anderson and Seton Lake black kokanee were most closely related to Portage Creek sockeye salmon, as previously found (Moreira & Taylor 2015), and kokanee from these two lakes were differentiable from each other based on outlier loci. Within the Columbia River drainage, the close relationships between kokanee (Okanagan, Wood, and Skaha Lakes) and anadromous sockeye salmon in the Okanagan River basin indicate these populations evolved from a common source and/or that there has been ongoing gene flow. The close connection between Skaha Lake kokanee, Okanagan Lake kokanee and Okanagan River sockeye salmon exemplifies these patterns. Skaha Lake kokanee spawn in the channel between Okanagan Lake and Skaha Lake, and Okanagan River sockeye salmon historically spawned in the same channel, before becoming isolated following construction of McIntyre Dam in 1916 (Veale & Russello 2016). In contrast, Kootenay Lake kokanee populations were highly divergent from the nearest populations sampled in the Columbia River drainage, more similar to the Anderson and Seton Lake kokanee. Taylor *et al*. (1996) similarly found kokanee in Kootenay Lake and nearby lakes to have a closer affinity to the Fraser River drainage rather than populations sampled in the Columbia River drainage, possibly due to changes in river connection; our results based on the large SNP dataset are consistent with this hypothesis.

At a finer level, shore- and stream-spawning kokanee ecotypes in each lake are sister taxa, highlighting the recurrent evolution of reproductive ecotypes. In all but one lake, kokanee ecotypes were genetically distinct from each other. The only shore- and stream-spawning *O. nerka* that could not be genetically differentiated by neutral and/or outlier loci were in Tchesinkut Lake, consistent with the findings of Frazer & Russello (2013). The vast majority of individuals spawn on the shores of an island in Tchesinkut Lake, where the “shore-spawners” were originally sampled in 2010 (Frazer & Russello 2013). The “stream-spawners” were sampled at an outflow and Drew Creek, however, the latter spawning activity only occurred in this small creek once Ministry personnel cleared the mouth allowing kokanee access (Frazer & Russello 2013). These results further suggest there may not be distinct ecotype populations in this lake, though additional molecular and behavioral studies will be necessary to investigate this to rule out recent divergence or on-going gene flow.

### Genomic bases for ecotype divergence

The majority of divergence between populations was between different catchments and regions, rather than between ecotypes. While some of this differentiation may be the result of divergent selection, it is likely that genetic drift between these isolated populations is primarily responsible for this pattern, especially given the lower estimated effective population sizes in some locations (Table 1). We did, however, identify a large number of highly differentiated genomic regions that may be under divergent selection between ecotype pairs, with some islands of divergence and outlier loci observed in multiple pairwise ecotype comparisons. We found three kinds of common patterns, each showing a potentially different basis for parallel evolution. First, we identified common outlier loci (e.g. R68810, R40949, R122749) with patterns of differentiation that were shared between ecotype pairs, both in terms of extent and direction. This pattern is suggestive of genetic variation that was present in ancestral populations that has then been divergently selected independently in multiple instances under similar selective regimes, or that have spread between adapted populations through introgression, potentially via other populations (Welch & Jiggins 2014). Second, we found genomic regions in many kokanee-sockeye salmon pairs where an outlier SNP or island of divergence were highlighted, but in each comparison, the exact SNPs showing differentiation were different. This indicates a gene(s) in this region may be repeatedly differentially selected between ecotypes through novel beneficial mutations occurring in each instance, or that novel neutral SNPs have arisen in each population linked to pre-existing variants that have since become targets for selection. Third, we observed many SNPs that were strongly differentiated in a single paired ecotype sampling, but not in others. While some of these may be false positives, others may be linked to genes that have been under directional selection specific to a given lake system or may be linked to polygenic traits, where mutations in completely different genes are able to contribute to a similar final phenotype. As with studies of this nature, further validation is required in order to provide more direct evidence that outlier behaviour reflects signatures of selection as well as to make any firm conclusions on the nature of that selection.

### Anadromous sockeye/kokanee outlier loci

While the ‘kokanee phenotype’ can be defined as a migratory ecotype, it constitutes a range of morphological, physiological and behavioral differences compared to anadromous sockeye salmon; therefore understanding the effects of genes linked to this phenotype is challenging. The outlier loci described in this study provide clues as to some of the genes potentially underlying these traits, though further research is required to ascertain the specific changes linked to each SNP, and how these genes affect phenotype. Many of the commonly shared outlier SNPs lie within, or adjacent to, genes with functions that could be related to adaptive traits. Here, we briefly discuss outlier loci that exhibited the greatest differentiation for which we have reliable annotations and that were partially validated through their shared detection across historically-isolated drainages (Table 3). There were, however hundreds of outlier SNPs detected that may also be worthy of further investigation. While we describe the annotated or nearest gene(s) for each of these SNPs, it remains possible that other nearby genes or regulatory sequences are the actual targets of selection.

R122749 contained the most commonly shared sockeye/kokanee outlier SNP, detected in five population comparisons and in both major river catchments. In both sockeye salmon populations, the C allele at this SNP was found at >90% frequency in each population, while for all kokanee populations the C allele was found at <10% frequency. R122749 maps to the angiopoetin-4 gene that regulates angiogenesis and the development of perivascular cells, along with regulating adipocyte differentiation and metabolism (Zhu *et al*. 2012). This gene is differentially selected and up-regulated in Tibetan chickens as a probable adaptation to hypoxia (Zhang *et al*. 2016), and dissolved O_2_ could constitute a target for differential selective pressure between kokanee and anadromous sockeye salmon given the variation in the environments they inhabit throughout their life cycle.

R40949 contained an outlier SNP in four kokanee/sockeye salmon comparisons, and is located in HSP90, a heat shock gene that regulates a variety of cellular processes in response to environmental temperature (Pratt & Toft 2003). Sockeye salmon populations display differences in thermal tolerance over both broad and fine spatial scales, and thermal stress has been identified as a major selective pressure (Eliason *et al*. 2011). Since kokanee and sockeye salmon experience greatly differing thermal regimes throughout their lifecycle, this gene constitutes a strong candidate to be differentially selected. QTL studies in Arctic char (*Salvelinus alpinus*) have previously demonstrated HSP90 divergence between thermal-tolerant and thermal-sensitive individuals (Quinn *et al*. 2011). Similarly, divergent selection at HSP90 has been demonstrated in sympatric *Anolis* lizards that occupy different thermal environments (Akashi *et al*. 2016). Interestingly, ubiquitin, which is often involved in the same pathways as and interacts with HSP90, has also been linked to thermal tolerance in Arctic char (Quinn *et al*. 2011), and ubiquitin conjugating enzyme was one of the other strong outliers recorded between Anderson/Seton Lake kokanee and Portage Creek sockeye salmon (R111962).

R7641 contained an outlier SNP in four kokanee/sockeye salmon comparisons and annotates to the Rac GTPase Activating Protein 1 (RACGAP1) gene. RACGAP1 has been shown to regulate sperm maturation in rainbow trout (Sambroni *et al*. 2013) and to be involved in complex signaling pathways that respond to varying water quality (Gagné & Cyr 2014).

R47083 is located in the dendritic cell-specific transmembrane protein (DC-STAMP) and was significant in three kokanee/sockeye salmon comparisons and one shore/stream comparison. This protein is related to bone re-development and haematopoesis regulation, along with having functions in immune homeostasis (Kobayashi *et al*. 2016). Variants in the DC-STAMP gene have been linked to size variation in humans, due to its regulation of bone remodeling activity (van der Valk *et al*. 2015). Given the different morphology and size of kokanee compared to anadromous sockeye salmon, this may be a candidate for divergent selection.

R18232 contained an outlier SNP that was significant in three kokanee/sockeye salmon comparisons and maps to the hepatocyte growth factor (HGF) gene, which regulates embryogenesis, organogenesis, and tissue regeneration by regulating cell growth and morphogenesis, and plays a central role in angiogenesis (Fajardo-Puerta *et al*. 2016). Previous QTL studies have highlighted this gene as being associated with altering growth rates in turbot (*Scophthalmus maximus*) (Robledo *et al*. 2016).

There are many other SNPs identified in this study that are worthy of further investigation. For instance, a total of 24 RAD tags (R7641, R11041, 13215, R14428, R18232, R22663, R24343, R40494, R45590, R46876, R47083, R50861, R59110, R61789, R68424, R68810, R74283, R80957, R89478, R93027, R96627, R106956, R114019, R122749) contained outliers independently in at least three population comparisons. Of these, R59110 was also recorded as an outlier between *O. nerka* ecotypes in Nichols *et al*. (2016), and maps to growth differentiation factor 3 (GDF3), another gene with strong potential for divergent selection between kokanee and anadromous sockeye salmon given its important role in ocular and skeletal development (Ye *et al*. 2010).

We also note that outlier loci not shared between comparisons may likewise be of interest, as they may be linked to novel mutations responsible for the complex and divergent kokanee and anadromous sockeye salmon phenotypes. For instance, R3833, R62994, R22554 and R37537 are all extremely highly differentiated between Anderson Lake and Seton Lake black kokanee and Portage Creek sockeye salmon. The unusual spawning behaviour and selective environment associated with the ‘black kokanee’ phenotype may potentially be linked with changes in the regions near these loci. Of particular note, R62994 was also the most significant outlier in Nichols *et al*. (2016). All outlier loci are given in Table S4, along with their alignments to the *Salmo salar* genome where possible.

**Table 3.** Annotations for high-confidence outlier loci independently detected across the Columbia and Fraser River catchments.

| RAD Tag | SNP | Accession No. | Gene | E-value |
| --- | --- | --- | --- | --- |
| 7641 | 37 | NC_027312.1 | RACGAP1 Rac GTPase activating protein 1 | 5.00E-37 |
| 18232 | 53 | NC_027315.1 | LOC106573447 hepatocyte growth factor-like | 1.00E-38 |
| 40949 | 10 | NC_027305.1 | LOC106608136 heat shock protein HSP 90-alpha-like | 4.00E-32 |
| 47083 | 30 | NC_027304.1 | LOC106605382 dendritic cell-specific transmembrane protein-like | 1.00E-39 |
| 68810 | 51 | NC_027308.1 | LOC106610979 leucine-rich repeat-containing protein 9-like | 2.00E-42 |
| 122749 | 24 | NC_027302.1 | LOC106598978 angiopoietin-related protein 4-like | 1.00E-38 |

### Islands of divergence

Outlier SNPs detected within one pairwise comparison and located adjacent to each other on the sockeye salmon linkage map are potentially closely physically linked, and thus may constitute islands of divergence exhibiting the same signal of differential selection.

Our formal tests for islands of divergence identified a number of potential genomic regions with increased divergence. While we used identical methods and linkage maps to Larson *et al*. (2017), the number of loci retained for our analyses was significantly fewer, as not all of our loci were able to be mapped unambiguously to their linkage map. The number of loci genotyped may affect the ability to identify islands of different sizes, with denser sequencing tending to reveal more and smaller islands (Nosil *et al*. 2012; Soria-Carrasco *et al*. 2014). Nevertheless, in the three primary comparisons, we described 12, 11 and 8 islands, a similar number to the 10 identified by Larson *et al*. (2017). In our study, the number of SNPs per island was on average slightly lower, and the percentage of outliers in the islands of divergence was likewise lower. This may be a consequence of the lower SNP coverage, differing scopes of the studies, and/or varying levels of population differentiation present in our respective systems.

Both the similarities and differences in location of islands of divergence between our results and those of Larson *et al*. (2017) are of interest and may show various patterns of divergent evolution or rapid genetic drift. We detected no evidence of divergence at the large island on linkage group 13 near the TULP4 gene, indicating that perhaps this is a novel mutation in the anadromous sockeye salmon in Alaska, rather than a more universal region underlying ecotype differentiation. We did, however, find some overlap in islands of divergence on linkage groups 12 and 15, which may represent regions that are rapidly evolving in each population, or are more generally under divergent selection among ecotypes. While there are many potential mechanisms that can create islands of divergence, when these are driven by adaptive divergence, strong selection and high gene flow is hypothesized to create fewer islands that vary in size, but display high differentiation. In contrast, weaker selection and low gene flow is hypothesized to create numerous small islands displaying lower differentiation (Nosil *et al*. 2009; Via & West 2008). In the study of beach- vs. stream- vs. river-spawning anadromous sockeye salmon in Alaska (Larson *et al*. 2017), few small islands of divergence with generally low divergences were found, suggesting high selective pressure acting on small segments of the genome under high gene flow. Our results showed similar numbers and sizes of islands of divergence, but with higher divergence outside these areas. While important differences in both the geographic scopes and density of SNPs between our study and that of Larson *et al*. (2017) may help account for these patterns, it seems likely that there is far greater isolation between ecotypes in our BC systems, leading to the effects of genetic drift being more prominent across the genome.

Genomic regions containing multiple outlier SNPs, but from different pairwise comparisons, also require further investigation to infer underlying mechanisms. Two of these regions identified particularly stand out: one on sockeye salmon linkage group 4a contained three outlier loci (R48535, R54026, R20531) all adjacent, but identified from three different kokanee/sockeye salmon comparisons; the other, on sockeye salmon linkage group 15a, also contained three outlier loci (R93027, R18325, R45460) that were identified from three kokanee/sockeye salmon comparisons and one kokanee shore/stream comparison. This pattern may be due to parallel directional selection on genes in this region, but in each population, novel mutations could either be responsible for, or linked to, these putatively adaptive changes. Alternatively, these regions may signal areas of the genome generally undergoing rapid selection within each population - similar to that experienced by immune genes like the major histocompatibility complex (MHC) (Larson *et al*. 2014a). The two known MHC gene regions for sockeye salmon are also located on linkage group 15a, though they are approximately 12 cM away from the region identified in our study according to the linkage map of Larson *et al*. (2016).

### The ‘shore-stream’ locus SNP 68810

The extremely high differentiation of the outlier SNP in R68810 between shore-spawning kokanee and stream-spawning kokanee or sockeye salmon in both the Columbia River and Fraser River drainages highlights the probable importance of a gene in this region influencing ecotype divergence. This locus was also identified as a high-confidence outlier by Nichols *et al*. (2016) in a comparison between shore-spawning sockeye salmon and stream-spawning kokanee in the Snake River catchment, further validating its potential significance. The only shore- /stream-spawning ecotype pair that did not show any divergence at this SNP was in Tchesinkut Lake, and as discussed above, there is no genetic evidence associated with ecotype differentiation in this location. In the West Arm of Kootenay Lake, while patterns of variation at this SNP were recorded in the same direction as observed in other comparisons (i.e. the G allele with higher prevalence in the shore-spawning population), differentiation was less significant. In this system, shore-spawning kokanee spawn on beaches near the sampled river mouths, and it remains unclear the degree to which these shore-and stream-spawning populations are reproductively isolated (Lemay & Russello 2012).

R68810 was not able to be mapped to the sockeye salmon linkage map (Larson *et al*. 2016) and was not assessed in the ecotype differentiation study of Larson *et al*. (2017). We were, however, able to map it to the leucine-rich repeat-containing protein-9 gene (LRRC9) in both the rainbow trout and Atlantic salmon genomes (*O. mykiss* chromosome 29, *S. salar* chromosome 9). LRRC proteins are involved in gene expression and participate in many biologically important processes, such as enzyme inhibition, hormone–receptor interactions, cell adhesion and cellular trafficking (Linhoff *et al*. 2001) A QTL-based study of mirror carp (*Cyprinus carpio* L.) hypothesized that LRRC proteins play an important role regulating the expression of superoxide dismutase (Xu *et al*. 2013), a commonly employed biomarker of immunotoxicity in different fish species (e.g. *Oryzias latipes*; Zelikoff 1998). There are 17 other genes within 250kb on either side of the LRRC9 gene in the *S. salar* genome, and it remains possible that the genetic variation under divergent selection is outside this area. Alternatively, the *O. nerka* genome may have structural changes compared to Atlantic salmon bringing other genes closer. We are currently investigating the generality of this locus as an ecotype identifier across the range of *O. nerka*, as well as the surrounding genomic regions for the linked genes under divergent selection.

Given the ecological, economic and cultural importance of kokanee in British Columbia, this highly informative SNP in combination with other outlier loci identified here have immediate utility for informing fisheries management throughout the province (Russello *et al*. 2012; Veale & Russello 2016) and likely across the entire range. We have already shown the efficacy of the five kokanee/sockeye salmon outliers (R7641, R18232, R40949, R47083, R122749) for informing a restocking program and associated spawning habitat management, through studies of hybridization rates between kokanee and introduced anadromous sockeye salmon in Skaha Lake, British Columbia (Veale & Russello 2016). The resolution for describing introgression levels obtained from the five outlier loci far surpassed the signal provided by 50 neutral genotyping assays (Veale & Russello 2016). Similarly, our results indicate that, in at least Okanagan Lake, shore- and stream-spawning kokanee can be identified to reproductive ecotype with high accuracy using a single genotyping assay, something not previously possible even with thousands of neutral markers (Lemay & Russello 2015). Moving forward, the genomic resources generated here hold great promise for establishing SNP panels that can accurately, rapidly and cost-effectively inform stock assessment, restocking and other fisheries management applications at multiple spatial scales, from the lake-level to cross-drainage programs and international initiatives.

## Acknowledgements

Richard Bussanich and Paul Askey contributed to initial project development. Rick Taylor and Amanda Moreira kindly provided samples. Matthew Lemay and Evelyn Jensen offered valuable feedback on the manuscript. Evelyn Jensen assisted in constructing the map figure and Eileen Klatt kindly provided permission to reproduce her sockeye salmon painting in several of the figures. Jim Seeb, Wesley Larson and Ryan Waples provided information and advice on analyses. Genome BC (award no.: UPP003), Okanagan Aquatic Enterprises, BC Hydro, and BC Ministry of Forests, Lands and Natural Resource provided funding to MR supported this research.

## Authors’ Contributions

AV carried out the molecular lab work and data analyses, participated in the design of the study and drafted the manuscript; MR conceived, designed and coordinated the study, assisted in data analyses and helped draft the manuscript.

